# Long-read transcriptome data of bee fungal parasite, *Nosema ceranae*

**DOI:** 10.1101/2020.03.11.987271

**Authors:** Huazhi Chen, Yu Du, Yuanchan Fan, Haibin Jiang, Cuiling Xiong, Yanzhen Zheng, Dafu Chen, Rui Guo

## Abstract

*Nosema ceranae*, a widespread fungal parasite that infects honeybee and many other bee species, can seriously affect bee health and colony productivity. In this article, *N. ceranae* spores were purified followed by third-generation sequencing using Nanopore PromethION platform. Totally, 6988795 raw reads were yielded from purified spores, with a length distribution among 1 kb~10 kb and a quality (Q) score distribution among Q6~Q12. A total of 6953469 clean reads were obtained, and among them 73.98% were identified as being full-length. The length of redundant reads-removed full-length transcripts was ranged from 1 kb to 5 kb, with the most abundant length of 1 kb. These data will improve transcriptome quality of *N. ceranae* significantly.

## Value of the data

- The dataset reported here allows a deeper understanding of the complexity of *Nosema ceranae* transcriptome.
- The accessible data can benefit researchers in identification of genes associated with *N. ceranae* infection mechanism.
- This dataset could be used to improve functional annotations of *N. ceranae* genome and transcriptome.
- The deposited data contributes to further exploring alternative splicing and polyadenylation of *N. ceranae* mRNAs.

## Data

The shared long-read transcriptome data were derived from purified *N. ceranae* spores. Based on Nanopore sequencing, 6988795 raw reads were produced, with an average length of 881 bp and a mean N50 of 971 bp (**Table 1**). As **Fig. 1** shown, these raw reads were ranged from 1 kb to 10 kb in length, with the most abundant length of 1 kb. Additionally, the quality (Q) score of majority of raw reads was Q9 (**Fig. 2**). A total of 6953469 clean reads were obtained, and among them 73.98% were identified as being full-length (**Table 2; Fig. 3**). After removing redundant reads, the length of remaining full-length transcripts was distributed among 1 kb~5 kb, with the highest percentage of 1 kb (**Fig. 4**).

**Table 1.** Summary of Nanopore raw reads

| Sample | Read num | Base num | N50 | Mean length | Maximun length |
| --- | --- | --- | --- | --- | --- |
| Nc | 6988795 | 6158102087 | 971 | 881 bp | 96051 bp |

**Table 2.** Summary of full-length clean reads

| Sample | Number of<br>clean reads | Number of<br>full-length clean reads | Percentage<br>of<br>full-length clean reads |
| --- | --- | --- | --- |
| Nc | 6953469 | 5143999 | 73.98% |

**Fig. 1.**
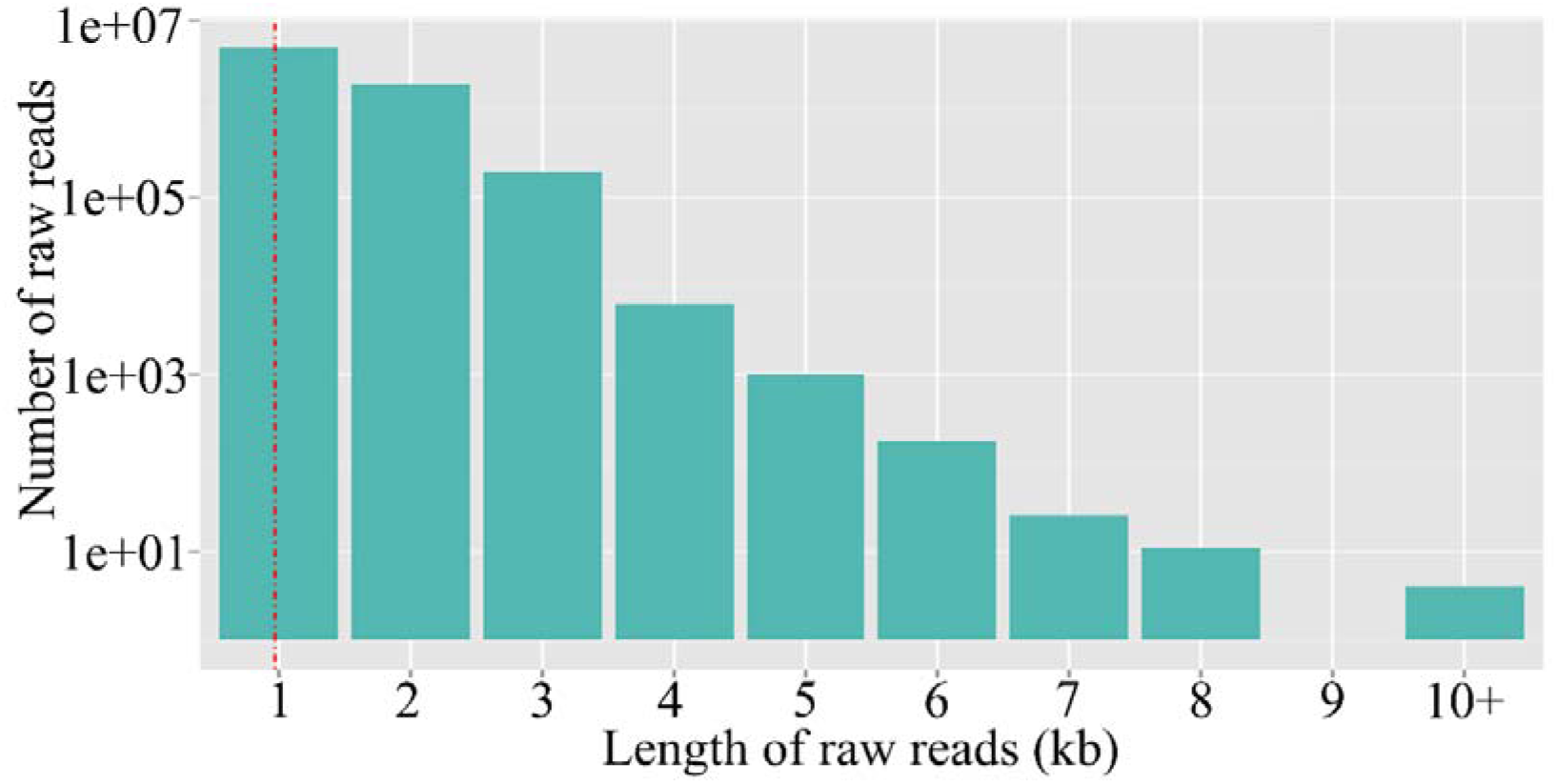
Length distribution of clean reads

**Fig. 2.**
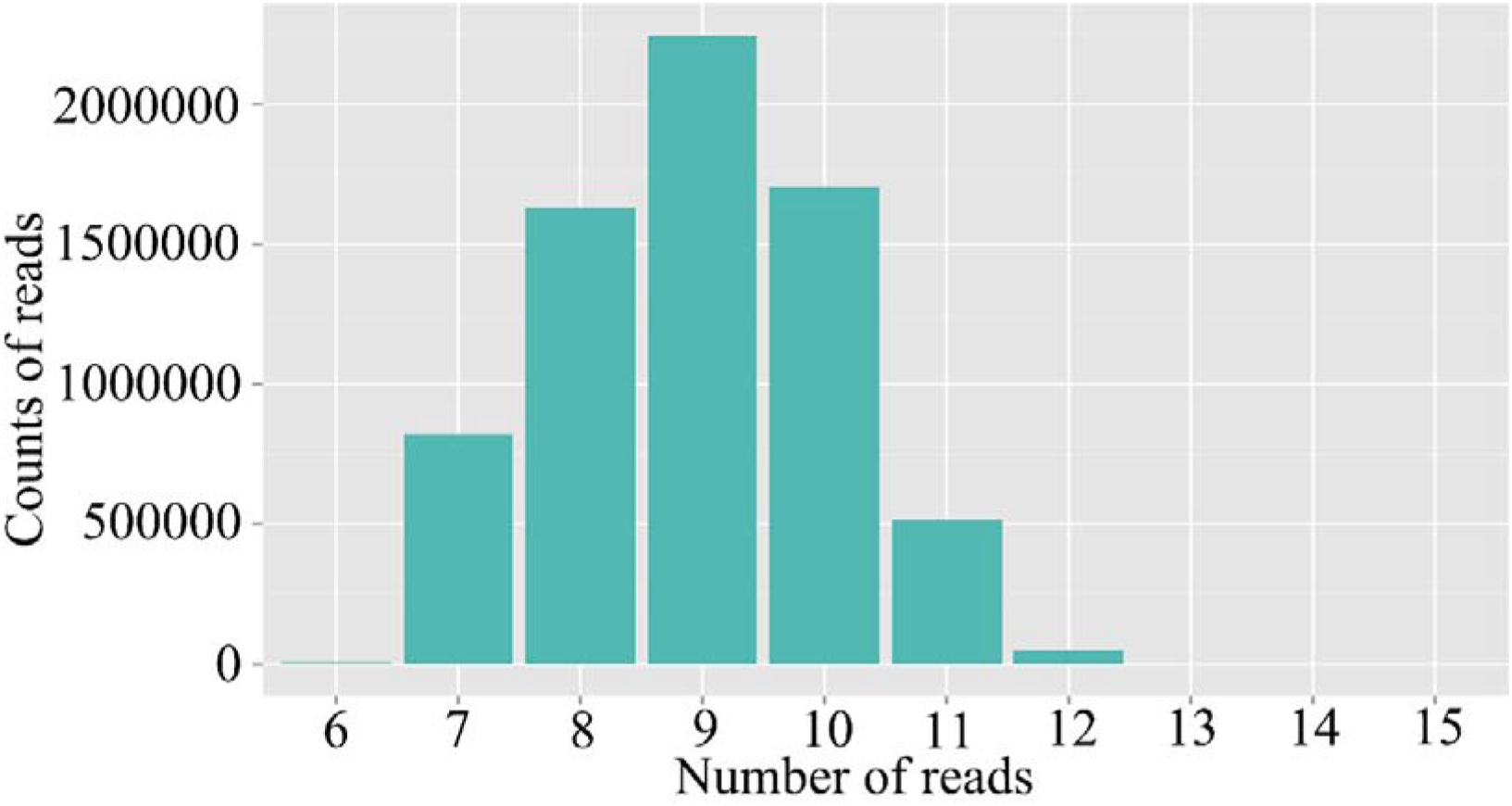
Quality distribution of clean reads

**Fig. 3.**
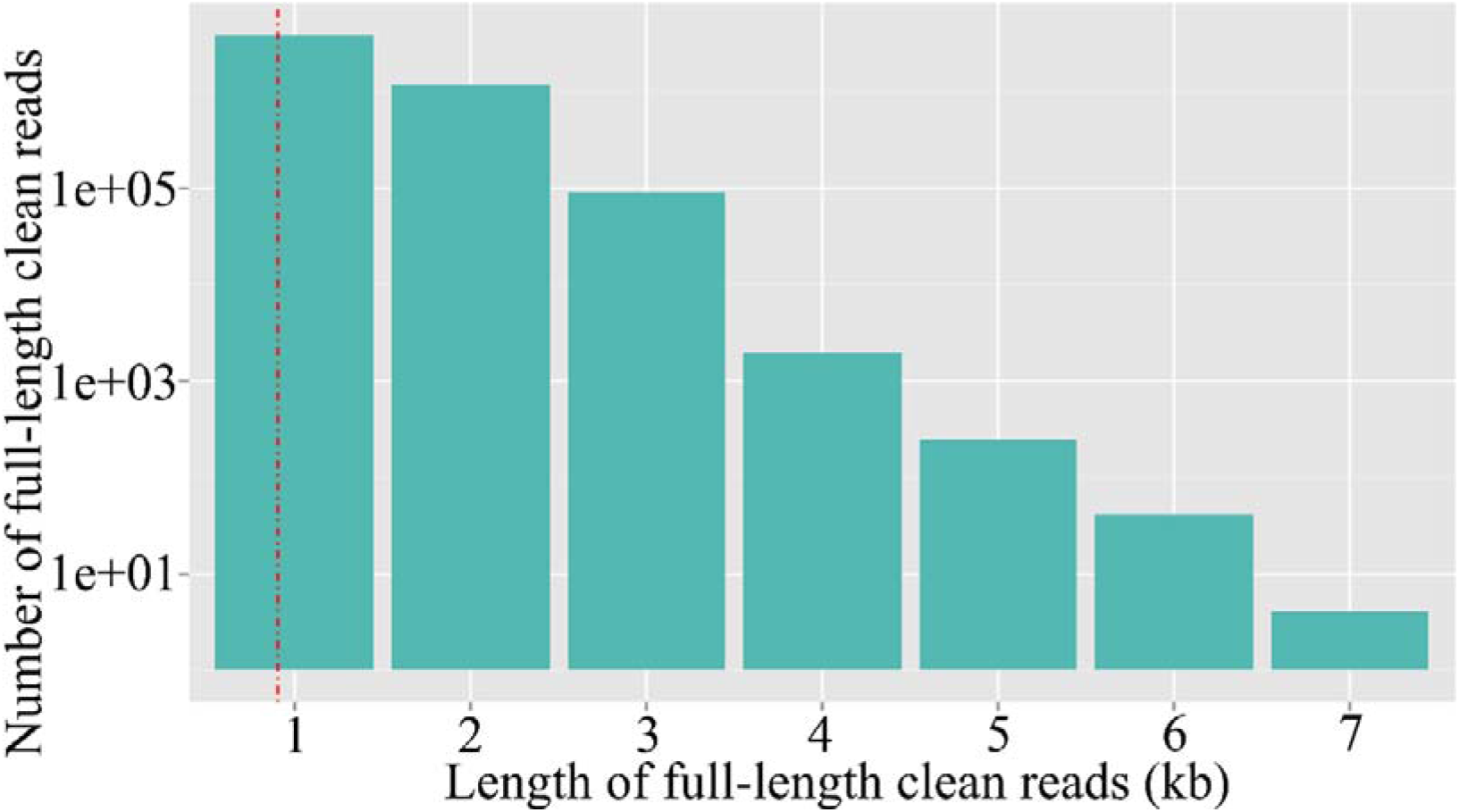
Length distribution of full-length clean reads

**Fig. 4.**
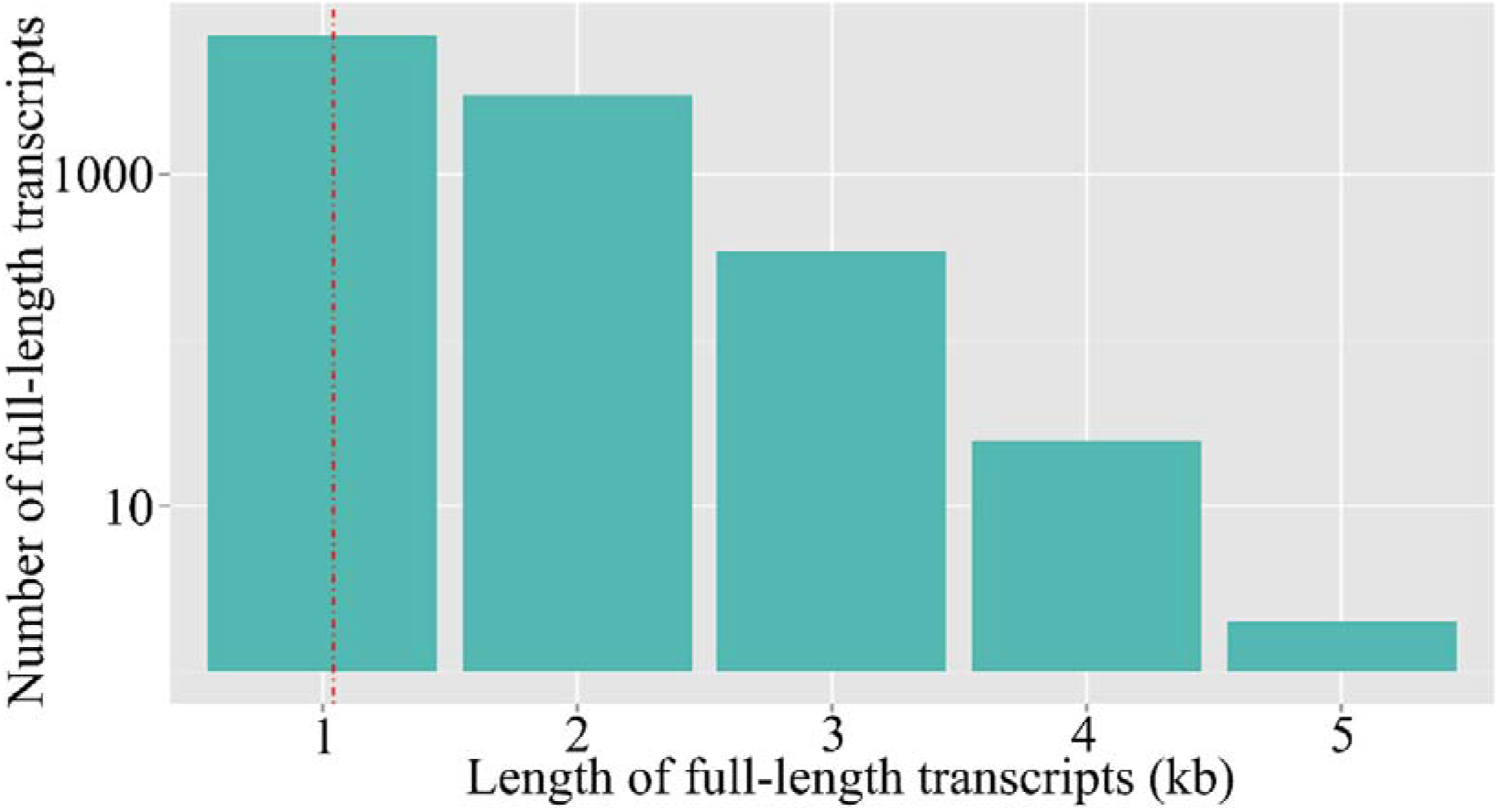
Length distribution of redundant reads-removed full-length transcripts

## Experimental Design, Materials, and Methods

### 2.1 Purification of N. ceranae spores

*N. ceranae* spores were previously purified following the method described by Cornman et al. [1] with some minor modifications [2]. A bit of spores was subjected to PCR detection and proved to be mono-specific [3] using previously developed specific primers [4]. The purified spores were immediately frozen in liquid nitrogen and stored at − 80 °C until use.

### 2.2 RNA isolation, cDNA library construction and Nanopore sequencing

Total RNA of *N. ceranae* spores was isolated using TRizol reagent (Thermo Fisher, China) on dry ice followed by reverse transcription with Maxima H Minus Reverse Transcriptase, and then purified and concentrated on a AMPure XP beads. cDNA libraries were constructed from 50 ng total RNA with a 14 cycles PCR using the cDNA-PCR Sequencing Kit (SQK-PCS109) and PCR Barcoding Kit (SQK-PBK004) according to the manufacturer’s protocol (Oxford Nanopore Technologies Ltd, UK), followed by sequencing by Biomarker Technologies (China) using PromethION platform (Oxford Nanopore Technologies Ltd, UK).

### 2.3 Long-read processing

Firstly, raw reads were filtered with minimum average read Q score=7 and minimum read length=500bp; ribosomal RNA were discarded after mapping to rRNA database. Secondly, full-length non-chemiric (FLNC) transcripts were determined by searching for primer at both ends of reads. Thirdly, clusters of FLNC transcripts were obtained after mapping to reference genome of *N. ceranae* (assembly ASM98816v1) with mimimap2 [5], and consensus isoforms were obtained after polishing within each cluster by pinfish (https://github.com/nanoporetech/pinfish); consensus sequences were mapped to reference genome (assembly ASM98816v1) using minimap2. Fourthly, mapped reads were further collapsed by cDNA_Cupcake package (https://github.com/Magdoll/cDNA_Cupcake) with min-coverage=85% and min-identity=90%; 5’ difference was not considered when collapsing redundant transcripts.

## Acknowledgments

This research was supported by the Earmarked Fund for China Agriculture Research System (No. CARS-44-KXJ7), the Science and Technology Planning Project of Fujian Province (No. 2018J05042), the Teaching and Scientific Research Fund of Education Department of Fujian Province (No. JAT170158), the Outstanding Scientific Research Manpower Fund of Fujian Agriculture and Forestry University (No. xjq201814), and the Scientific and Technical Innovation Fund of Fujian Agriculture and Forestry University (No. CXZX2017342, No. CXZX2017343).

